# Analysis-ready datasets for insecticide resistance phenotype and genotype frequency in African malaria vectors

**DOI:** 10.1101/582510

**Authors:** Catherine L. Moyes, Antoinette Wiebe, Katherine Gleave, Anna Trett, Penelope A. Hancock, Germain Gil Padonou, Mouhamadou S. Chouaïbou, Arthur Sovi, Sara A. Abuelmaali, Eric Ochomo, Christophe Antonio-Nkondjio, Dereje Dengela, Hitoshi Kawada, Roch K. Dabire, Martin J. Donnelly, Charles Mbogo, Christen Fornadel, Michael Coleman

## Abstract

The impact of insecticide resistance in malaria vectors is poorly understood and quantified. Here a series of geospatial datasets for insecticide resistance in malaria vectors are provided so that trends in resistance in time and space can be quantified and the impact of resistance found in wild populations on malaria transmission in Africa can be assessed. Data are also provided for common genetic markers of resistance to support analyses of whether these genetic data can improve the ability to monitor resistance in low resource settings. Specifically, data have been collated and geopositioned for the prevalence of insecticide resistance, as measured by standard bioassays, in representative samples of individual species or species complexes. Data are provided for the *Anopheles gambiae* species complex, the *Anopheles funestus* subgroup, and for nine individual vector species. In addition, allele frequencies for known resistance associated markers in the Voltage-gated sodium channel (*Vgsc*) are provided. In total, eight analysis-ready, standardised, geopositioned datasets encompassing over 20,000 African mosquito collections between 1957 and 2017 are provided.

## Background & Summary

Current malaria control activities are heavily reliant on vector control using insecticides, which means resistance to these compounds has the potential to derail control efforts ^1,2^. Studies have started to investigate the impact of resistance in certain situations ^3,4^ but a full understanding of impact requires comprehensive quantification of resistance. To quantify the factors that influence vector control, data from vector populations are required and a number of vector databases are already available for species distributions, infection prevalence, and bionomic parameters ^5–12^. A database for insecticide resistance in malaria vectors, that allows users to download analysis-ready datasets, is vital so that the impact of levels of resistance found in wild populations on malaria transmission can be assessed. These datasets are also essential to quantify trends in resistance in space and time, filling the gaps in the available data with robust predictions, to aid resistance management and the deployment of interventions designed to counter resistance ^13^.

Studies of phenotypes in natural populations may be confounded by variation in the environments sampled, including factors linked to climate, land use and malaria control interventions. It is not possible to control for all variables in the natural environment but this issue can, in part, be mitigated by sampling a large number of locations encompassing different combinations of environmental variables. Large, collated datasets do, however, have potential disadvantages. Collated datasets that are a combination of data points representing different types of sample, different measurement methods, different location types and so on, risk undermining any analysis that is performed ^6,14^.

Each dataset should be constructed to address a specific question or set of questions, and the data within each set needs to be standardised to allow robust analyses. The goal of the current work was to collate data from multiple studies characterising the insecticide resistance phenotype and genotype in communities of malaria vectors at as many locations and times as possible. The aim was then to generate standardised datasets designed to address specific questions using geospatial analyses. Namely, what are the trends in resistance in time and space in specific vector assemblages, to then assess whether these are associated with trends in malaria transmission. The second aim was to provide data that can be used to investigate associations between genetic markers for individual mechanisms of resistance and the insecticide resistance phenotype, to assess whether genetic markers can improve the ability to monitor resistance in low resource settings ^15^.

## Methods

### Data sources

Published articles were identified in the Web of Science bibliographic database by using the search terms “insecticide resistance” and “anopheles” together with the name of each malaria endemic country in turn. The Web of Science was chosen because it incorporates many relevant databases including the SciELO Citation Index from 1997 onwards, MEDLINE from 1950 onwards (from the U.S. Library of Medicine), the Data Citation Index from 1993 onwards (provides details of datasets in international data depositories), the BIOSIS Citation Index from 1969 onwards (covers pre-clinical, experimental, and animal research) and the Web of Science’s own Core Collection from 1945 to date.

The earliest date was unrestricted, and the search was completed on 31 December 2017. The initial search yielded 3,685 articles published from 1956 to 2017, with the first African paper published in 1957. Data were extracted from each article as outlined below and 342 articles provided data from field samples of mosquitoes collected in malaria endemic African countries for either the insecticide resistance phenotype and/or genotype. If values for some data fields were missing, the authors were contacted. In these instances, either i) the phenotype/genotype data was given in the article but supplementary information such as the date of sampling or mosquito identification method was missing, or ii) the genotype/phenotype data were missing, or had been aggregated across sites or years, so the disaggregated data for each site-year were requested. In the latter instance, any genotype/phenotype data received from the authors were treated as unpublished. In addition, groups undertaking vector surveillance, or involved in large studies that had not yet published their results, were asked to provide unpublished datasets. In total, 42 unpublished datasets from African countries were provided.

### Data aggregation / disaggregation

The aim of this work was to provide measures of insecticide resistance for representative samples of a species population (or a species complex or a subgroup) found at a particular time and place, rather than data at the level of an individual mosquito. Replicates from the same mosquito collection sampled at a single “site” and “collection period” were aggregated. The spatial resolution of a “site” was defined by the original field studies and classified by the current study, as described in the data geo-referencing section below. The temporal resolution of a “collection period” was also defined by the original data generators and the duration of each collection period was recorded in the current dataset, as described below. If the reported data had been pooled across multiple sites or collection periods, but was originally obtained at a finer resolution, the disaggregated data for each site-period were requested. For example, if mosquitoes were collected from five sites and bioassayed separately giving a bioassay result for each site, but only a single average result for the region was published, then the five separate results were requested.

Datasets were constructed based on mosquito samples that represented either a single species or a species complex or subgroup. Species-level data were entered wherever it was available and aggregated to provide data for the species complex later, using the original species composition. If insecticide resistance data were provided for each species but the original species composition was not available for that study, the data points for each individual species were included in species-level datasets (provided they met the inclusion criteria below) but they were not aggregated to provide data for the species complex.

### Inclusion criteria

Subgroup-, complex- or species-level insecticide resistance phenotype data generated from either a WHO susceptibility test ^16–26^ or a CDC bottle bioassay ^27^ using either the F0 or F1 generation from a field collection of *Anopheles* mosquitoes were included. Data were excluded if the susceptible strain control failed, i.e. mortality in the susceptible strain was <100%.

Subgroup-, complex- or species-level data on the resistance variants in the voltage-gated sodium channel (*Vgsc*) gene that were derived from F0 or F1 generations from field collections of *Anopheles* mosquitoes, provided as either genotype or allele frequencies, were included.

Only mosquito samples that were representative of a species complex or subgroup and/or a species were included and any samples that were subject to sub-setting that biased the original sample were excluded. For example, if a mixed species sample was collected but a bioassay result was only reported for the most common species, that bioassay result cannot be considered as representative of the species complex at that time and place. In this example, the data were included in the species-level datasets released here, but no values were included in the datasets for species complexes. Similarly, if a mixed species sample was collected and then the F1 generation was sorted into single species by identifying the mother of each egg batch, those results cannot be considered representative of the species complex at that time and place. If the allele frequency was calculated for mosquitoes that survived a bioassay, and dead mosquitoes were not tested, this result cannot be considered as representative for either the species complex, or the individual species, and was not included in any dataset. If a mixture of dead and alive mosquitoes from a bioassay were tested to obtain an allele frequency, but the ratio of dead:alive was not representative of the original sample, for example 80% died in the bioassay but the sample tested was 50:50, then these data were also excluded.

### Individual data files

The full database was used to generate eight individual data files (Table 1) that address specific questions for defined sets of mosquitoes. The aim in creating data files 1 and 2 was to provide a set of comparable results for each insecticide from bioassays that had used the same insecticide concentration and exposure duration, however, the recommended concentrations and durations varied with WHO protocol version. The protocol version used to define the standard insecticide concentrations and exposure durations in data file 1 and 2 was the 1998 WHO test procedures, because the highest volumes of data across all years were available for the concentrations and durations specified by this protocol version ^25^. Insecticides that were not covered by the 1998 protocol version were specified in the 2013 version so this later protocol was also used to set the standard values for data files 1 and 2 ^26^.

**Table 1.**
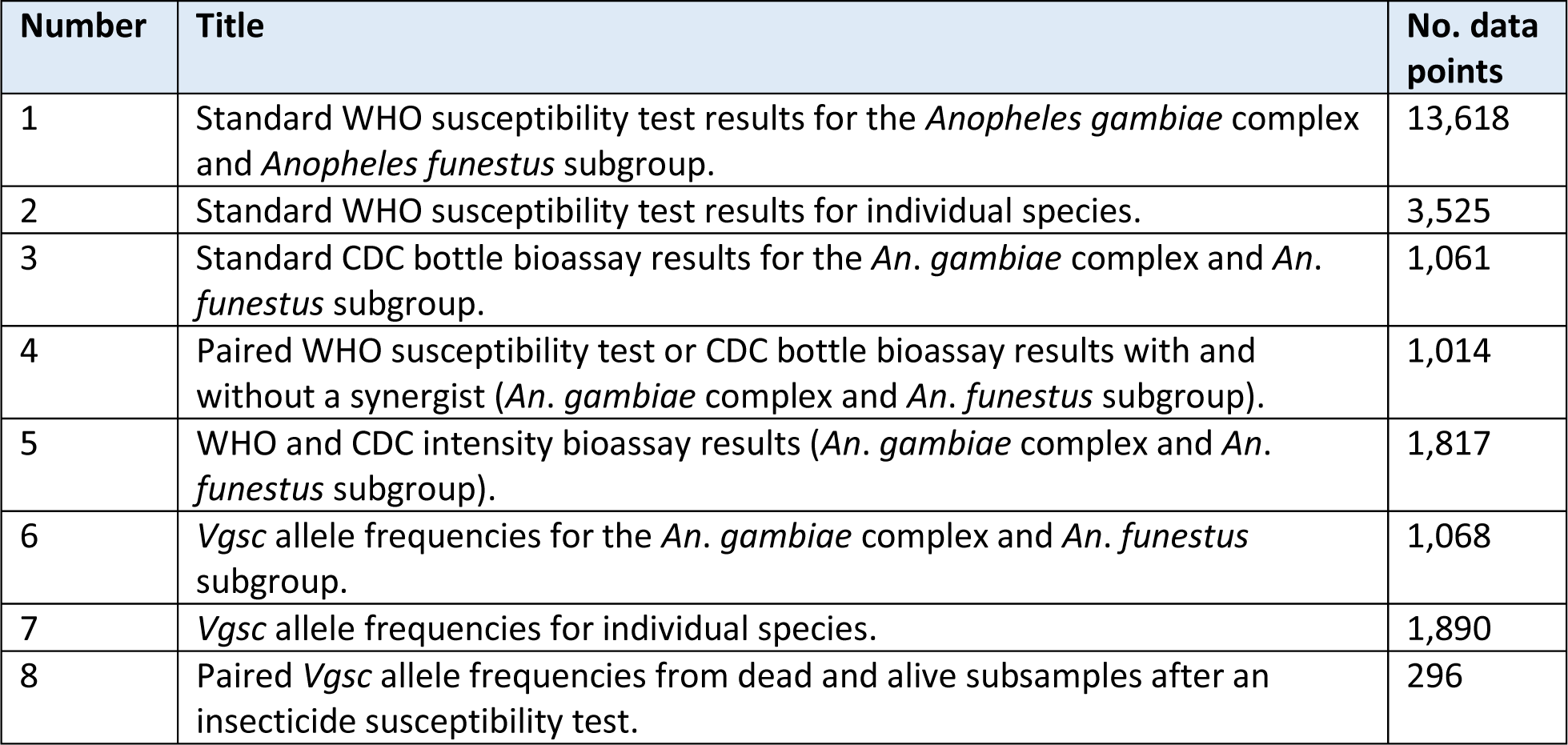
Summary of each of the eight data files released.

### Data fields

The data fields included in this release are described in Tables 2–7. The source data fields (Table 2), the sample collection data fields (Table 3), and the geo-location data fields (Table 4) are provided in all data files. The species identification data fields (Table 5) are provided in data files 2,4,7,8. The bioassay data fields (Table 6) are provided in data files 1,2,3,4,5. The *Vgsc* data fields (Table 7) are provided in data files 6,7,8.

**Table 2.** Data source data fields.

| Title | Data type | Description |
| --- | --- | --- |
| Source citation | Text | Citation for the data source. This field is duplicated up to four times to record instances where the full information linked to that data point came from more than one source. |
| Source type | Category | Each source is categorised as 'journal article', 'personal communication' and so on. |

**Table 3.** Sample collection data fields.

| Title | Data type | Description |
| --- | --- | --- |
| Capture method | Category | The method used to capture the mosquito sample. This field is duplicated four times to record instances where samples from different capture methods were pooled before testing. |
| Start month | Integer | The month when the mosquito collection began. |
| Start year | Integer | The year when the mosquito collection began. |
| End month | Integer | The month when the mosquito collection ended. |
| End year | Integer | The year when the mosquito collection ended. |

**Table 4.** Geo-locations data fields. These data fields are described further in the data geo-referencing section of the text.

| Title | Data type | Description |
| --- | --- | --- |
| Country | Category | The country that the site sampled was in. |
| Site type | Category | Sites can be a point, a polygon or multiple-points, as described in the geo-positioning section. |
| Site name | Free text | Name of the field site sampled. |
| Latitude | Decimal number | For 'point' sites only, the geographical coordinates are given in decimal degrees. This field is duplicated multiple times to record instances where samples from more than one 'point' site were pooled before testing. |
| Longitude | Decimal number | For 'point' sites only, the geographical coordinates are given in decimal degrees. This field is duplicated multiple times to record instances where samples from more than one 'point' site were pooled before testing. |
| Admin level | Category | If the site is a 'polygon' that matches an administrative unit, the administrative level (0, 1 or 2) is recorded. |
| GAUL code | Integer | If the site is a 'polygon' that matches an administrative unit, the identifier from the Global Administrative Units Layer is recorded. |

**Table 5.** Species identification data fields.

| Title | Data type | Description |
| --- | --- | --- |
| Identification method | Category | The molecular method used to identify individual species in the original sample. This field is duplicated to record instances where two methods were used. |

**Table 6.** WHO and CDC bioassay data fields.

| Title | Data type | Description |
| --- | --- | --- |
| Anopheline | Category | The species or species complex or subgroup that the bioassay result represents. |
| Generation | Category | The mosquito generation tested: F0, F1 or a mix of both. |
| Test protocol | Category | The WHO or CDC bioassay protocol followed is listed. |
| Insecticide | Category | The insecticide tested is named. |
| Concentration (%) | Decimal number | If a WHO protocol was followed, the insecticide concentration is given as a percent. |
| Concentration (µg/bottle) | Decimal number | If the CDC protocol was followed, the insecticide concentration is given in µg/bottle. |
| Exposure period (minutes) | Integer | The period of exposure to the insecticide in minutes. |
| No. mosquitoes tested | Integer | The total number of mosquitoes tested in all replicates. |
| No. mosquitoes dead | Integer | The total number of mosquitoes that died in all replicates. |
| Percent mortality | Decimal number | The percent of mosquitoes that died across all replicates, adjusted using Abbot's formula if applicable. |

**Table 7.** *Vgsc* gene data fields.

| Title | Data type | Description |
| --- | --- | --- |
| Anophelines tested | Category | The species or species complex or subgroup that the genetic result represents. |
| Method | Category | The molecular method used to identify alleles. This field is duplicated three times to record instances where up to three different methods were used on the same sample. |
| Generation | Category | The mosquito generation tested: F0, F1 or a mix of both. |
| No. mosquitoes tested | Integer | The total number of mosquitoes tested. |
| **Genotype frequencies** |  |  |
| L/L (no.) | Integer | The number of mosquitoes homozygous for the wildtype, susceptible allele (1014L). |
| L/L (%) | Decimal number | The percent of mosquitoes homozygous for the wildtype, susceptible allele (1014L). |
| L/F (no.) | Integer | The number of mosquitoes heterozygous for the wildtype, susceptible allele (1014L) and the 1014F resistance allele. |
| L/F (%) | Decimal number | The percent of mosquitoes heterozygous for the wildtype, susceptible allele (1014L) and the 1014F resistance allele. |
| L/S (no.) | Integer | The number of mosquitoes heterozygous for the wildtype, susceptible allele (1014L) and the 1014S resistance allele. |
| L/S (%) | Decimal number | The percent of mosquitoes heterozygous for the wildtype, susceptible allele (1014L) and the 1014S resistance allele. |
| F/F (no.) | Integer | The number of mosquitoes homozygous for the 1014F resistance allele. |
| F/F (%) | Decimal number | The percent of mosquitoes homozygous for the 1014F resistance allele. |
| F/S (no.) | Integer | The number of mosquitoes heterozygous for the 1014F and 1014S resistance alleles. |
| F/S (%) | Decimal number | The percent of mosquitoes heterozygous for the 1014F and 1014S resistance alleles. |
| S/S (no.) | Integer | The number of mosquitoes homozygous for the 1014S resistance allele. |
| S/S (%) | Decimal number | The percent of mosquitoes homozygous for the 1014S resistance allele. |
| **Allele frequencies** |  |  |
| L1014L | Decimal number | The allele frequency (%) for the 1014L wildtype, susceptible allele. |
| L1014F | Decimal number | The allele frequency (%) for the 1014F resistance allele. |
| L1014S | Decimal number | The allele frequency (%) for the 1014s resistance allele. |

All of the data fields were extracted and entered as they were provided by the source. If any information was missing, no value was entered (see the missing data section below) and the authors were contacted. The only values that were generated after the data were extracted from the sources, were the geo-location values. Full details on how the geo-location data were generated is given in the next section.

Data files are provided for species complexes or subgroups, and for individual species, separately. Bioassay data for individual species were obtained from studies that either sorted egg batches based on mothers’ species prior to a bioassay being performed, or disaggregated the results by species after the bioassay was performed, or instances where all mosquitoes in the original sample were found to be one species after the bioassay was performed.

For data files 4, an additional identifier (the “matched set ID”) is included to allow results from bioassays that used the same mosquito collection and exactly the same bioassay conditions, with or without a synergist, to be identified. The same approach was used for data files 5 where the “matched set ID” allows results from bioassays that used the same mosquito collection and exactly the same bioassay conditions, but with differing insecticide concentrations and/or exposure durations, to be identified. In total, 453 matched sets are provided in the synergist dataset, 464 in the intensity assay data file, and 148 for *Vgsc* allele frequencies in paired dead and alive subsamples.

### Data geo-referencing

In order to use these data in geospatial models at a resolution of ∼5km, each mosquito collection location was classed as either a point (defined as a site located within a 2.5 arc-minute grid cell, i.e. an area of ∼5 × ∼5 km) or a polygon (defined as a site with an area greater than that of a point). For all sites defined as ‘points’ the following steps were followed. The site name and all contextual information about the location of the site were noted, for example, the district the site was in, its proximity to a major city or other geographical feature, and so on. If the data source provided coordinates, then these were converted to decimal degrees. If no coordinates were provided the site name was searched in at least two online gazetteers (Google Maps, GeoNames, OpenStreetMap, WikiMapia and so on). All options identified by this search were cross-checked against the contextual information. If only one option matched the contextual information, the coordinates were extracted from the online gazeteer and added to the database. If more than one option matched the contextual information, or no options were found that matched the contextual information, the individuals who published or provided the data were contacted. In these instances, no coordinates were entered without external confirmation. After all possible coordinates were obtained for a study, they were plotted on a map to ensure the data spread for that study matched any information available on the authors’ overarching sampling strategy.

For all sites defined as ‘polygons’, any contextual information was noted, such as the province that the district was in. The name of the area in question was searched in the FAO’s Global Administrative Unit Layers (GAUL, http://www.fao.org/geonetwork/srv/en/metadata), using fuzzy matching to allow for different spellings or transliteration, and checked against any available contextual information. If one administrative unit in GAUL matched the area name and contextual information, the GAUL code (= a unique identifier for that area/polygon) was extracted and entered in the database. If an administrative unit within GAUL could not be identified, no code was entered.

If an individual site could not be located, or could not be precisely located within a 2.5 arc-minute grid cell, then the data point was linked to the second order administrative division that the site falls within. The administrative division was identified using the same method as for polygons above.

If multiple point locations were sampled and the mosquitoes were pooled before being tested (or only the pooled results were available), the site type was classified as a ‘multi-point’ and the coordinates for all of the individual point locations were linked to the test result.

### Missing data

If data for a particular field was missing from the original data source, the value was recorded as NR, i.e. not reported. For values that were not applicable, rather than missing, NA was used. For example, if there was only one data source linked to a data point, the value for the second data source was NA. If the geographical coordinates for a site could not be identified (see above), NF was entered, i.e. not found.

If a study did not explicitly state the insecticide concentration, exposure period and/or minimum number of mosquitoes used, but did specify the protocol followed, it may be possible to obtain the missing information from the relevant protocol ^16–27^. Protocol values for the most commonly used insecticides are provided in Tables 7 and 8, and the values for all insecticides are given in Supplementary Information file 1, to allow data users to fill these data gaps if they wish.

**Table 8.** Minimum recommended number of mosquitoes and duration of exposure specified by published protocols for the WHO susceptibility test and CDC bottle bioassay.

| Protocol | Min. no. mosquitoes | Duration (minutes)<sup>1</sup> | Duration for fenitrothion (minutes) | Duration for DDT (minutes) |
| --- | --- | --- | --- | --- |
| WHO 1963 | 60 | 60 |  | 60 |
| WHO 1970 | 60 | 60 | 60 | 60 |
| WHO 1975 | 60 | 60 | 60 | 60 |
| WHO 1976 | 60 | 60 | 60 | 60 |
| WHO 1980 | 60 | 60 | 120 | 60 |
| WHO 1981 | 60 | 60 | 120 | 60 |
| WHO 1986 | 60 | 60 | 120 | 60 |
| WHO 1992 | 60 | 60 | 120 | 60 |
| WHO 1998 | 80 | 60 | 120 | 60 |
| WHO 2013 | 80 | 60 | 120 | 60 |
| WHO 2016 | 80 | 60 | 120 | 60 |
| CDC bottle bioassay | 100 | 30 | 30 | 45 |

<sup>1</sup>The exposure duration values from the WHO protocols apply to dieldrin, malathion, fenthion, propoxur, chlorphoxim, permethrin, deltamethrin, $\lambda$ -cyhalothrin, bendiocarb, etofenprox, pirimiphos-methyl, carbosulfan, cyfluthrin, chlorfenapyr, fipronil and $\alpha$ -cypermethrin. The CDC protocol values apply to bendiocarb, cyfluthrin, cypermethrin, deltamethrin, fenitrothion, $\lambda$ -cyhalothrin, malathion, permethrin and pirimiphos-methyl. Full details can be found in the published protocols.

**Table 8.** Insecticide concentrations specified by published protocols for the WHO susceptibility test and CDC bottle bioassay. M.C. denotes that multiple concentrations were recommended so the actual concentration used in any particular bioassay cannot be inferred from the protocol version.

| Protocol | DDT | deltamethrin | permethrin | bendiocarb | λ-cyhalothrin | primiphos-methyl |
| --- | --- | --- | --- | --- | --- | --- |
| WHO 1963 | M.C. |  |  |  |  |  |
| WHO 1970 | M.C. |  |  |  |  |  |
| WHO 1975 | M.C. |  |  |  |  |  |
| WHO 1976 | M.C. |  |  |  |  |  |
| WHO 1980 | 4% | 0.025% | 0.25% |  |  |  |
| WHO 1981 | 4% | 0.025% | 0.25% | 0.1% |  |  |
| WHO 1986 | 4% | 0.025% | 0.25% | 0.1% |  |  |
| WHO 1992 | 4% | 0.025% | 0.25% | 0.1% | 0.1% |  |
| WHO 1998 | 4% | 0.05% | 0.25% | 0.1% | 0.05% |  |
| WHO 2013 | 4% | 0.05% | 0.25% | 0.1% | 0.05% | 0.25% |
| WHO 2016 | 4% | 0.05% | 0.25% | 0.1% | 0.05% | 0.25% |
| CDC bottle bioassay | 100 µg/bottle | 12.5 µg/bottle | 21.5 µg/bottle | 12.5 µg/bottle | 12.5 µg/bottle | 20 µg/bottle |

### Data duplication

The data extracted came from several hundred different sources, which introduced the possibility that individual results had been entered into the database more than once. To identify duplicates the following data fields were used: original sample; species tested; date fields; no. mosquitoes tested; no. mosquitoes dead; percent mortality; site name; coordinates. Fuzzy matching was used for all fields to identify duplicates where different levels of aggregation had been used, or different data values were missing, or names were spelled differently. All partial matches were examined to identify genuine duplicates. Duplicate data points were removed, and the source details linked to the single data point that was retained. In total, 3,483 duplicated data points were removed.

### Data ownership and permissions

Data ownership is retained by the original data owners. For all unpublished data, permission to include these data in this release was requested. Of a total of 11,057 unpublished data points, permission was received to release 10,834.

## Data Records

The data are available for download from the Dryad Digital Repository: https://doi.org/10.5061/dryad.dn4676s [link will work after acceptance of the manuscript]. The spatial and temporal distributions of data file 1, standard WHO susceptibility test results for the Anopheles gambiae complex and Anopheles funestus subgroup, are shown in Figure 1. The spatial and temporal distributions of data file 7, Vgsc allele frequencies for individual species, are shown in Figure 2.

**Figure 1.**
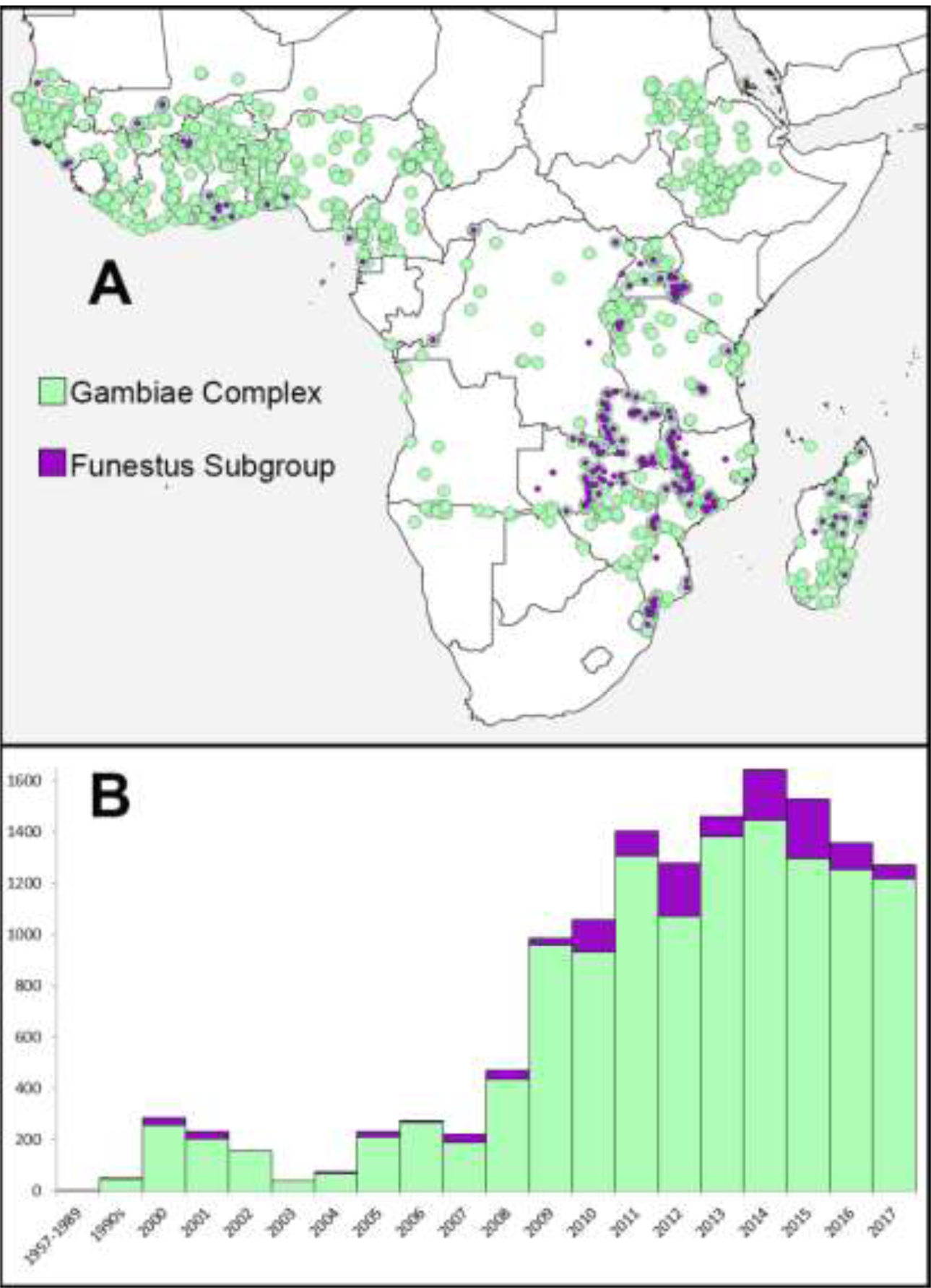
Spatial and temporal distributions of Data File 1. A. The locations of mosquito collections of the *An*. *gambiae* complex and the *An*. *funestus* subgroup that were used in standard WHO susceptibility tests. B. The number of data points available for each year for the *An*. *gambiae* complex (green) and the *An*. *funestus* subgroup (purple).

**Figure 2.**
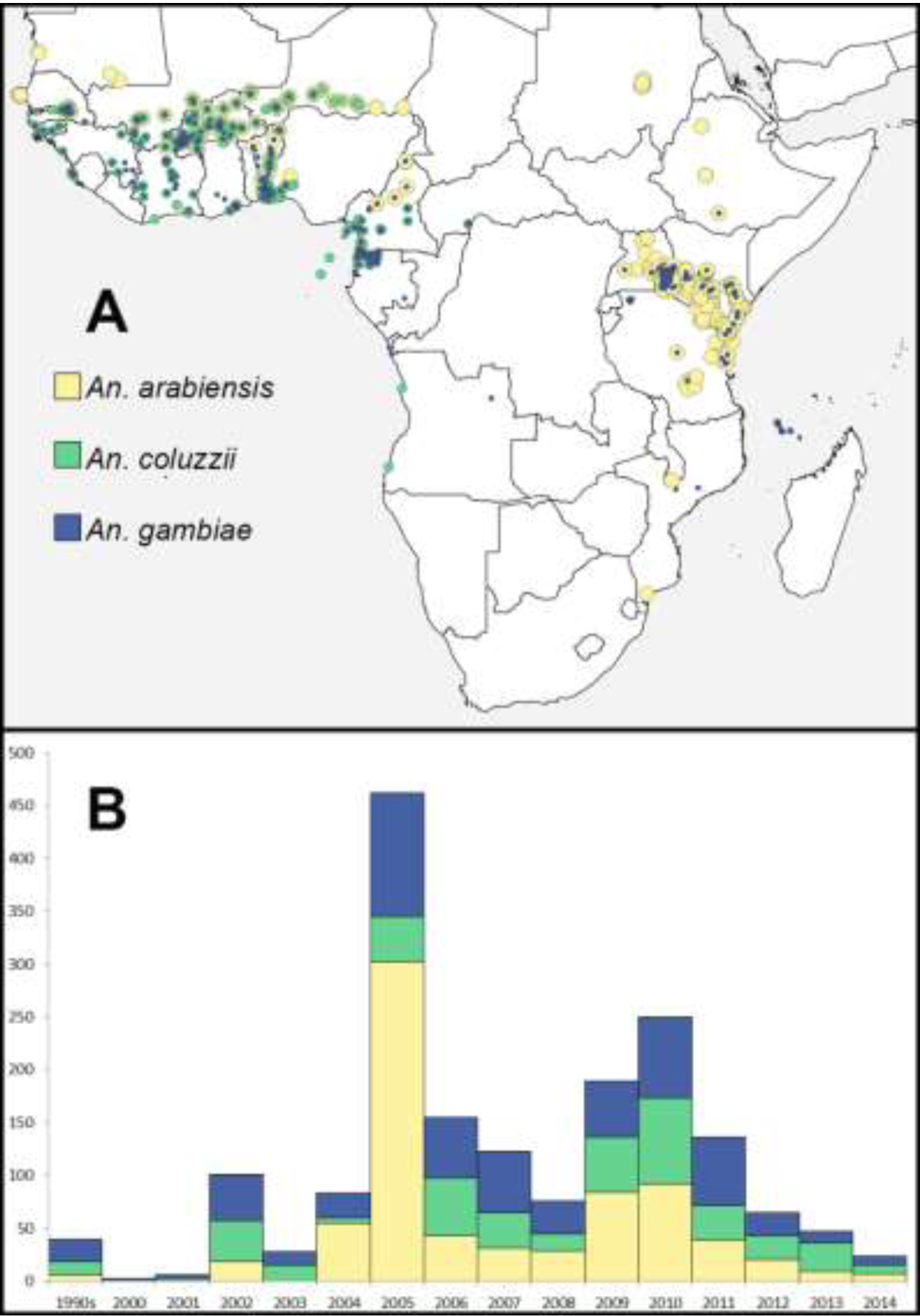
Spatial and temporal distributions of Data File 7. A. The locations of mosquito collections of *An*. *arabiensis, An*. *coluzzii* and *An*. *gambiae* that were used to calculate *Vgsc* allele frequencies. B. The number of data points available for each year.

Data file 1 is the largest dataset but all eight have similar spatial distributions with clustered sampling in the east and west of Africa and sparse data points in the centre and southwest. They also share similar temporal distributions with phenotypic data volumes increasing throughout the time period particularly from 2008 onwards, and the genotypic data volumes peaking in 2005 and 2010. The genotype data were almost exclusively extracted from published papers and there is typically a lag of around two years between mosquito collection and the publication of a paper containing the test results.

In addition to the data extracted for *Vgsc* allele frequencies, data were also identified for *Ace-1* allele frequencies and metabolic mechanisms of resistance including cytochrome P450s, esterases and glutathione-*S-*transferases. The volumes of genetic and biochemical data currently available for these mechanisms of resistance did not meet our aim of providing standardised data for a large number of locations across Africa, so no collated datasets for these mechanisms were generated. Dataset 4 consists of results from synergist bioassays so it does, therefore, provide data linked to P450-mediated mechanisms of resistance.

Many studies performed both bioassays and genetic tests. If links between the different tests performed on the same sample of mosquitoes were provided by the original study, and providing any subsamples tested were not biased, then it was possible to extract pairs of phenotypic and genotypic measures of resistance for samples from a specific time and place. Unfortunately, however, when instances of paired phenotypic and genotypic results for an individual species from a single time and place were extracted, only sixty pairs were identified. This volume of data did not meet our aim of providing standardised data for a large number of locations across Africa. The same was true for paired phenotypic and genotypic results for a species complex or subgroup from a single time and place.

In addition, the data volumes available for species-level CDC bottle bioassay results, species-level paired bioassays with and without a synergist, and species-level intensity bioassays, were too low to meet our aim of providing standardised data for a large number of locations across Africa.

## Technical Validation

Data were checked for internal consistency to ensure i) all coordinates for point locations fell on land and in the right country, as defined by GAUL, ii) mortality and allele frequencies never exceeded 100%, iii) the collection end date was never earlier than the collection start date, and iv) the species name tallied with the identification methods listed. A matrix of species identification methods and species identified by each method was prepared in order to complete this check (Supplementary Information file 2). In addition, a second person reviewed the geographical coordinates in accordance with the geo-locations protocol outlined above.

## Usage Notes

Each data file released has been designed to provide results for a representative sample of a species complex or subgroup, or an individual species, so users can be confident of what each set of results represents.

The data files have also been designed for use in geospatial analyses and, in such analyses, the precise location for each data point is important for two reasons. First, because this allows accurate calculation of the Euclidian distances between points for analyses that exploit spatial correlations in the data ^15,28^. Secondly, precise location information allows accurate matching of the data to a wide range of environmental variables, such as climatic, socio-economic and intervention variables, to exploit relationships between the biological data and these environmental variables ^29,30^. The use of data linked to wider areas is a current area of research aimed at improving model predictions in circumstances where data linked to precise locations are particularly sparse ^31,32^. For any kind of spatial analysis, it is essential to know whether the geographical coordinates provided represent a precise location or wider area, what the definition of a precise location is, and where the boundaries of the wider areas lie. The data points released here are linked to a mixture of precise locations and wider areas, the precise locations (referred to here as points) are defined as an area within a 2.5 arc-minute grid cell (approx. 5 × 5 km), and links to the boundaries of wider areas (referred to here as polygons) are given.

The data files released here are not the result of one single, continent-wide study that used a standard sampling design. It is a compilation of many studies that used many designs and incorporates obvious sampling bias. Sites that are more easily accessed or closer to research centres may be more likely to be sampled. Sites where high levels of resistance are expected may also be more likely to be sampled, as are sites where insecticide-based interventions are planned as a result of a combination of related variables. Geostatistical models can, however, be used to model sampling intensity to check these biases before proceeding further ^33^.

Other data resources for insecticide resistance in malaria vector are available. Data on insecticide resistance in the *Anopheles* vectors of malaria are also available from VectorBase, however, VectorBase’s aim and scope are much broader than those of the current data release and the data volumes for insecticide resistance in *Anopheles* vectors are smaller than those provided here ^34^. Furthermore, these data have not been configured specifically for use in mathematical analyses including geospatial analyses. Insecticide resistance data can also be viewed on interactive maps using the IR Mapper and Malaria Threats websites but these are data visualisation tools ^35^. The data shown on these sites were not collated in support of mathematical analyses and are not available for download. There are overlaps in all of these databases, including overlaps with the data being released here. The data released here includes data that were provided to the World Health Organization to support the establishment of the Malaria Threats website ^36^. In addition, the data being released here was shared with the group behind the IR Mapper site so that both groups could cross-check each other’s sources to identify publications that had been missed.

A geo-database of insecticide resistance in the *Aedes* vectors of arboviruses has previously been released but this has much smaller data volumes for Africa, and encompasses a much greater range of insecticides tested at a greater range of concentrations on both adults and larvae ^37^. In contrast, the data released here provide sufficient volumes of standardised values to support a range of analyses of insecticide resistance in malaria vectors in Africa and are freely available to all. In addition to the data release described here, these data have been shared with the Pan Africa Mosquito Control Association to support the establishment of an Africa -led and –managed data resource. The datasets released here will also be available from the IR Mapper and Malaria Atlas Project websites [www.irmapper.com; www.map.ox.ac.uk]. In addition, predicted values for the prevalence of resistance (i.e. mortality in a standard WHO susceptibility test) at every location in a ∼5km resolution grid, for each year from 2005 to 2017, will be modelled and released in the coming months.

## Data Citation

Moyes, C.L. *et al*. Data from: Analysis-ready datasets for insecticide resistance phenotype and genotype frequency in African malaria vectors. Dryad Digital Repository. doi:10.5061/dryad.dn4676s (2019).

## Acknowledgements

The preparation of this geospatial data resource was funded by the Wellcome Trust (grant 108440/Z/15/Z) and initial scoping work was funded by the Bill & Melinda Gates Foundation via the Vector-borne Disease Network (VecNet). U. S. President’s Malaria Initiative provided resistance data collected through PMI Africa Indoor Residual Project from several countries in Africa.

## Author information

### Contributions

C.L.M. devised the analysis-ready datasets for release. C.L.M. and A.W. drafted the data extraction and geopositioning protocols. A.W., K.G. and A.T. extracted, processed and geopositioned the data with guidance from C.L.M. and M.C. A.W. classified the species identification methods. K.G. extracted recommended sample sizes, doses and exposure durations from the WHO and CDC protocols. A.T. and A.W. identified duplicates in the data. C.L.M. checked the above work and reviewed the data. C.F., C.M., M.J.D., R.K.D., H.K., D.D., A.-N. C., E.O., S.A.A., A.S., C.S.M. and G.G.P. provided large volumes of unpublished data and provided advice on the use and format of these data. P.H., M.C. and C.L.M. contributed to the potential dataset uses.

### Competing interests

The authors declare no competing interests.

## Supplementary information

1. Insecticide concentrations stipulated by the published WHO insecticide susceptibility test and CDC bottle bioassay protocols.
2. Matrix of species identification methods.

